# BOTANIC-0: a series of foundation models for plant genomic data

**DOI:** 10.64898/2026.02.23.706817

**Authors:** Jean Ogier du Terrail, Tanguy Marchand, Vincent Cabeli, Zhor Khadir, Cyril Véran, Léonard Strouk

## Abstract

Genomic language models (gLMs) have emerged as a powerful paradigm for learning regulatory biology directly from DNA sequence. Here, we introduce `Botanic0`^¶^, a family of plant genomic foundation models spanning 100M to 1B parameters and pretrained on 43 phylogenetically diverse plant genomes. The `Botanic0`-S, `Botanic0`-M, and `Botanic0`-L models form the first generation of a long-term research initiative, dedicated to advancing crop improvement research, genotype–to-phenotype modeling, and sequence-based genome editing. The architecture, pre-training pipeline and pre-training dataset of `Botanic0` follow the seminal work of [1]. Across a broad suite of genomic and genetic prediction tasks, including regulatory element annotation, gene expression inference, and variant effect prediction, `Botanic0` models achieve performance competitive with state-of-the-art foundation models, both in zero-shot settings and after fine-tuning. Scaling analyses reveal consistent improvements in predictive power with increased model capacity, highlighting the benefits of large-model pretraining for plant genomics. This work establishes our ability to train foundation models at scale, and lays the foundation for the next generations of models to come. To support reproducible research and community benchmarking, we release all `Botanic0` models at https://huggingface.co/living-models/models.

## 1 Introduction

Accelerating climate change intensifies the race to develop resilient, high-yielding crops, posing a fundamental challenge to global food security. Yet the development cycle for improved cultivars remains slow: the timeline from trait discovery to varietal release spans many years—up to 8 years for a new variety [2]— while resistance genes are frequently overcome within just a few growing seasons as pathogens expand their geographic range and adapt to new environments [3]. As climate pressures intensify [4, 5] and yield gains lag behind projected global demand [6, 7], there is a critical need for approaches that can rapidly identify, prioritize, and manipulate the genetic variants shaping agronomically important traits.

Modern genomics has made it routine to map trait-associated regions through genome-wide association studies, pan-genomic analyses, and large-scale sequencing. However, translating these associations into mechanistic understanding remains one of the central bottlenecks in plant genetics. Determining which variants are causal, and how they influence gene regulation, cellular state, or organismal performance, still requires extensive experimentation, making it difficult to evaluate large numbers of candidate loci or to design genome edits with predictable phenotypic outcomes [8, 9]. The complexity and evolutionary diversity of plant genomes, including extensive noncoding regions, structural variation, and lineage-specific regulatory architectures, further complicates variant interpretation. Self-supervised deep learning, inspired by breakthroughs in natural language processing [10], provides a promising framework to decipher this genomic language.

Building on the success of large language models trained on natural text [11, 12, 13, 10, 14], genomic language models (gLMs) learn statistical and functional constraints directly from raw DNA sequences, without requiring labeled data [15]. Studies across domains have shown that such models capture sequence grammar, regulatory motifs, chromatin structure, and evolutionary constraints down to single-nucleotide resolution [16, 17, 18, 19, 20]. Foundational DNA models such as GENA-LM [21], GROVER [22], and Evo 2 [23] demonstrate that sufficiently large models trained on diverse genomes acquire generalizable representations that support a wide range of downstream prediction tasks.

In plants, this paradigm has only recently begun to take shape. To the best of our knowledge, GPN [19] was among the first works that demonstrated that pretraining convolutional neural networks (CNNs) with masked language modeling (MLM) allowed to build powerful plant gLMs as GPN trained on 8 Brassicales genomes yielded strong predictive performance when tested on a representative species from that clade, *Arabidopsis thaliana*. Building on this foundation, AgroNT [1], PlantCaduceus [24, 25], and related efforts [26] extended the approach by scaling gLM models, leveraging newer architectures [11, 16, 27], to diverse plant genomes spanning multiple orders. These studies showed that multispecies pretraining enables accurate prediction of regulatory features, tissue-specific gene expression, splice sites, and functional variants—even in species with sparse experimental annotations. Collectively, these works underscore the benefits of multispecies training, particularly in plants, where genome size, architecture, and repeat content vary dramatically across clades. By introducing new AI-ready plant benchmarks, including the Plant Genomic Benchmark (PGB) [1] and curated datasets from cross-species studies [24, 25], these efforts have fostered greater standardization in evaluation practices for plant gLMs, enabling subsequent work to build upon a shared and reproducible foundation.

Nonetheless, the intrinsic complexity of genomics—despite substantial community efforts to enhance reproducibility [28, 29, 30]—complicates the standardization of AI model evaluation in plant genomics. Moreover, the application of large-scale AI models to plant genomics remains relatively recent, and evaluation practices are still evolving. We anticipate that greater consensus will emerge as the field matures. To ensure fair comparisons and preserve methodological simplicity in this transitional phase, we therefore evaluate our models exclusively on the original, unmodified versions of these datasets.

In this report, we share our experience in setting up a MLM pre-training pipeline on plant genomics data loosely following the [1] pre-training pipeline. We generated three foundation models of different sizes `Botanic0-S` (114M parameters), `Botanic0`-M (260M parameters), and `Botanic0-L` (991M parameters) that were trained on a total of 43 species (cf. Section 4.1). We show that, similarly to previously cited works, our models can be used as natural embeddings for plant genomic-related machine learning tasks (cf. Section 2.3.1). `Botanic0` models can also be fine-tuned directly to specific tasks (cf. Section 2.4).

Finally, this work represents an initial step toward turning genomic foundation models into practical tools for crop improvement research, shortening the road from lab to field. Our scaling experiments (cf. Section 2.3.3) as well as learnings from the literature will be used to design the next generations of the `Botanic`-family models.

## 2 Results

In this section, we present the results of the experiments we conducted to evaluate the training dynamics as well as the final performance of the `Botanic0-S`, `Botanic0`-M, and `Botanic0-L` models and to compare them to the state-of-the-art.

While the masked language modeling (MLM) pre-training methodology we followed to build the `Botanic0` models is standard [1], implementation details, notably the specifics of dataset construction, preprocessing, and the training loop, are detailed in Section 4.1.

### 2.1 Pre-training Botanic0 models of different sizes at scale generalizes to unseen species

Figure 1 shows the 6-mer MLM cross-entropy loss evolution for the `Botanic0-S`, `Botanic0-M` and `Botanic0-L` models during the pre-training, which is done on 43 different species (see Section 4.1). We also display the estimation of the total pre-training loss on different checkpoints of different models on both the training set and on the out-of-distribution validation set composed of the 5 left out species on the same figure. We note that it is an estimation as the MLM batch construction process is inherently stochastic.

**Figure 1.**
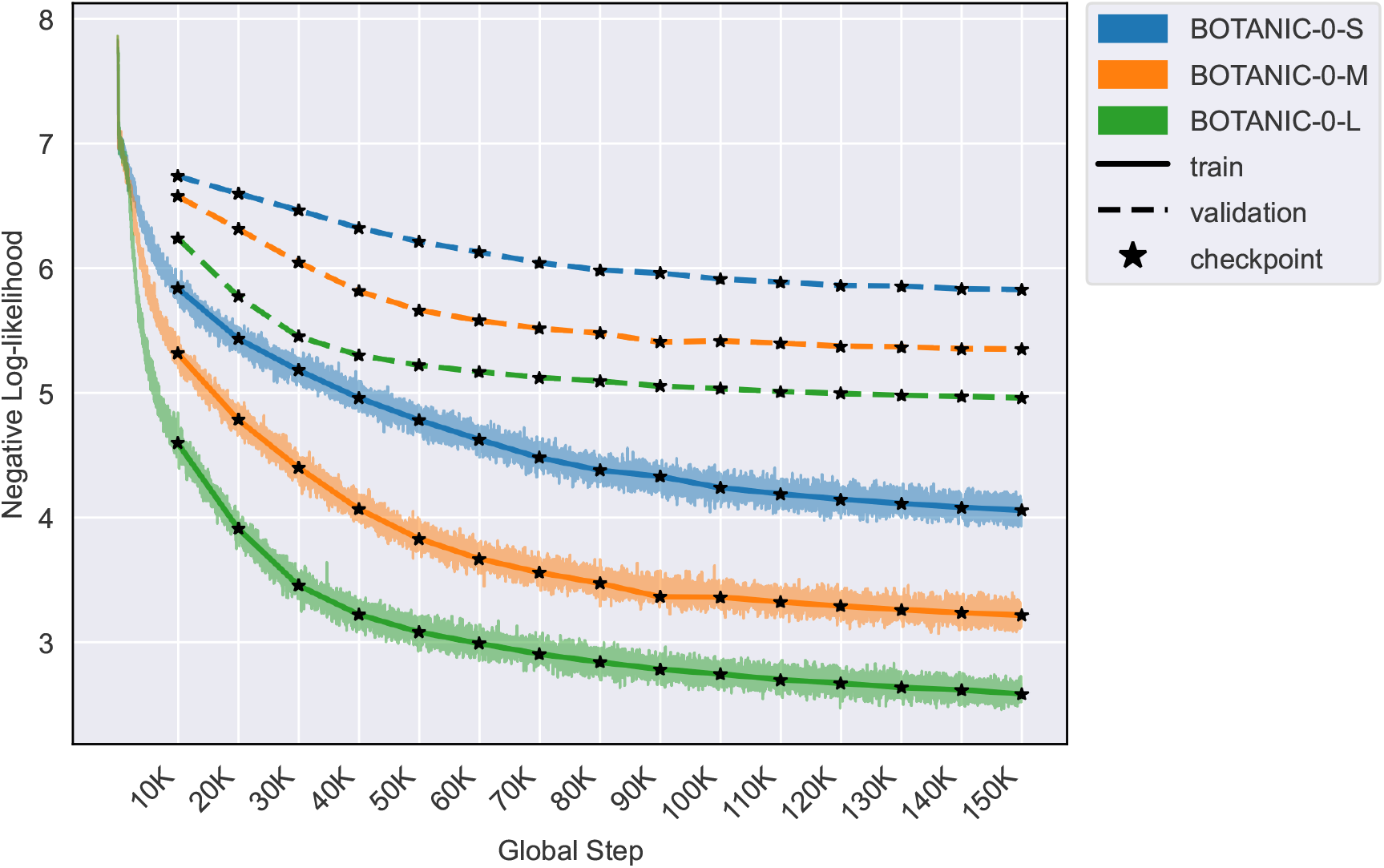
Pre-training loss analysis: per-step accumulated and synced across-workers aggregated loss, overlaid with total loss estimation on training and validation sets for a restricted set of checkpoints. Training dataset is comprised of 43 species, and validation dataset of 5 species unseen during training.

We observe that the pre-training loss is continuously decreasing over time for all models, indicating that the models are learning to predict masked 6-mers of DNA for species in the training set given a context of roughly 6000 base pairs. Furthermore, the validation loss is also decreasing over time for all models, indicating that the models are not overfitting to the training set and that transferability to unseen species is possible, although the validation loss is much higher than the training loss as shown in Figure 1. Most strikingly, we see no saturation of the validation losses with increasing model sizes, strongly suggesting that this approach is scalable to larger models.

### 2.2 Botanic0 zero-shot log-likelihood scores allow ranking deleterious mutations

After the pre-training, we computed the zero-shot log-likelihood ratio (LLR) between reference alleles and various deleterious mutations in *Arabidopsis thaliana*. The LLR, first introduced in [31], has been characterized in the literature as a good metric for functional constraint prediction: models that are successfully trained to understand DNA grammar will assign lower probabilities to deviations from the reference sequence that are likely to have a functional impact [15]. According to [24], both PlantCAD [24] and GPN [19] models are better at ranking deleterious mutations than AgroNT [1]. Our results show that `Botanic0-L` LLR scores achieve much higher correlation with those of PlantCAD and GPN models than AgroNT (Figure S4). More quantitative experiments would be needed to confirm whether or not this translates to better ranking performance.

### 2.3 Pre-trained Botanic0 embeddings extract biologically meaningful features

#### 2.3.1 Botanic0 embeddings capture elementary genomic structures

As a first step, we ensured that `Botanic0` embeddings were able to capture biologically meaningful features by performing a genomic region classification task in *Arabidopsis thaliana*. Figure 2a shows the confusion matrix of the genomic region classification task using an XGBoost classifier [32] trained on the last hidden state embeddings of the `Botanic0-L` model, and Figure 2b shows a visualization of the same embeddings projected into a 2D space using pymde [33]. We see qualitatively that distinct genomic regions are clustered into distinct regions of the space. Quantitatively the multi-class balanced accuracy of an XGBoost classifier trained on top of `Botanic0-` Lembeddings is 0.641 compared to 0.542 when trained on top of AgroNT [1] embeddings using the same procedure (see Supplementary Figure S3). Additionally, the CDS class splits into three distinct clusters in the embedding space, corresponding to the three codon reading-frame phases (see Supplementary Figure S1). It is likely that it is an artifact of the 6-mer tokenization. We add another view of the same plot in Supplementary Figure S2 to better visualize the ncRNA class as this class is not well separated from the others in the embedding space.

**Figure 2.**
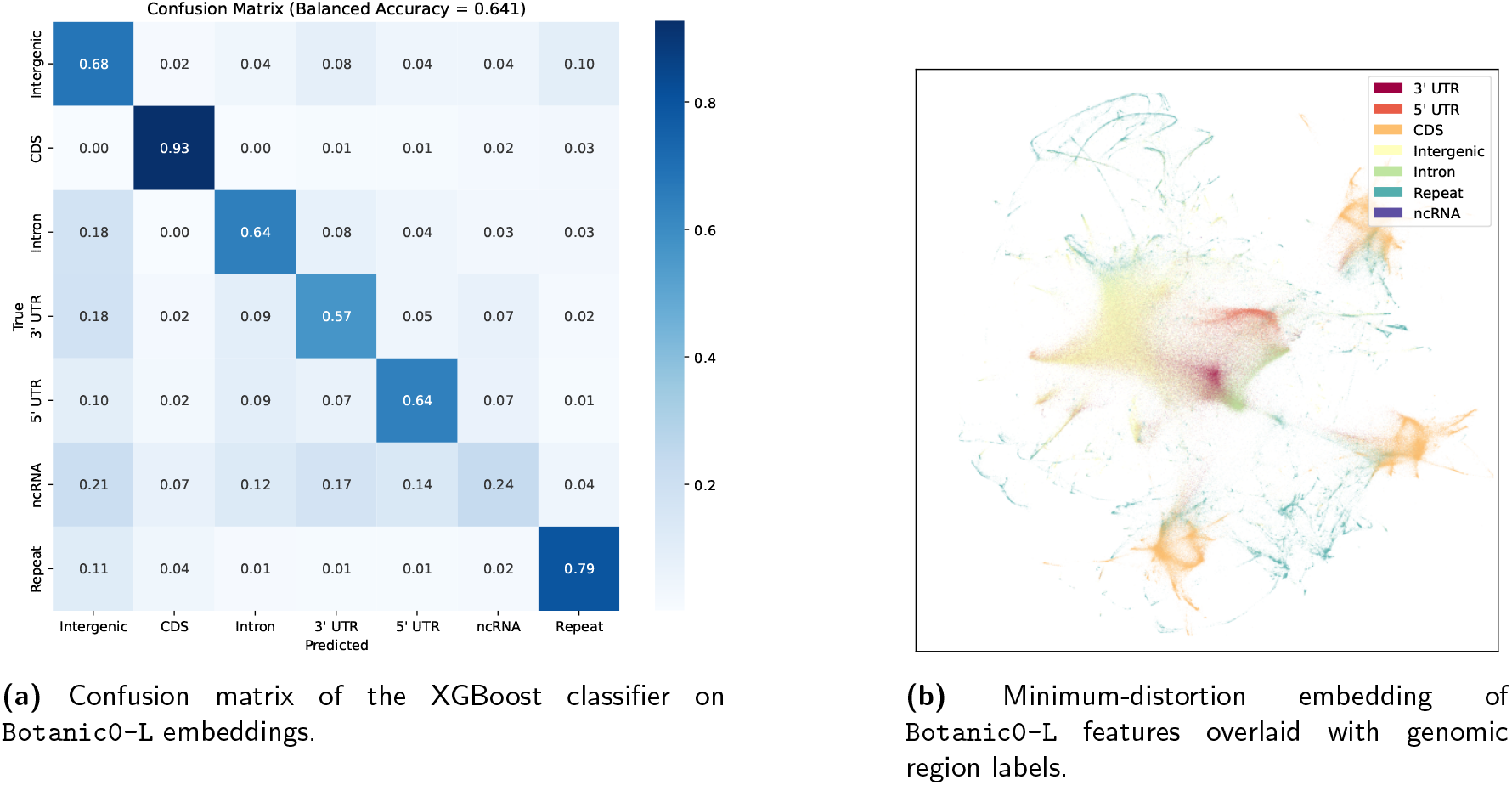
Genomic region classification in *Arabidopsis thaliana* from `Botanic0-L` embeddings. Cf Section 4.1.5 for technical details. The ncRNA class is diffuse in the embedding space therefore does not stand out in this plot. Better seen in color.

#### 2.3.2 Competitive performance of Botanic0 embeddings on standard benchmarks reflects their ability to capture biological information

In order to assess the quality of the learned representations we perform two kinds of probing tasks while keeping the weights of the models fixed. We use classic linear probe techniques as well as XGBoost probing following [24] on five binary classification tasks (PlantCAD-TIS, PlantCAD-TTS, PlantCAD-Donor, PlantCAD-Acceptor, PlantCAD-Conservation) from [24] (details are provided in section 4.2.3). Probing techniques are deemed more reliable than fine-tuning as a quality metric for comparing pretrained embeddings as fine-tuning entangles representation quality with the specifics of the optimization process [34].

The results, reported in Figure 3, show that `Botanic0` models are competitive with the best models in the field. Additionally, using the strand information available in four of the PlantCAD datasets to convert everything to (+) strand representation further improves the performance of models that are not RC-equivariant (all models except the PlantCAD family). The results also show that NTv2 [18] consistently underperforms on all tasks compared to all other competitor models, while its structure and size are close to other models’ in the benchmark (cf. Table 1). This shows that the pre-training dataset (NTv2 is trained on a mixture of plants and other living organisms) and the pre-training pipeline can have a dramatic impact on downstream tasks.

**Figure 3.**
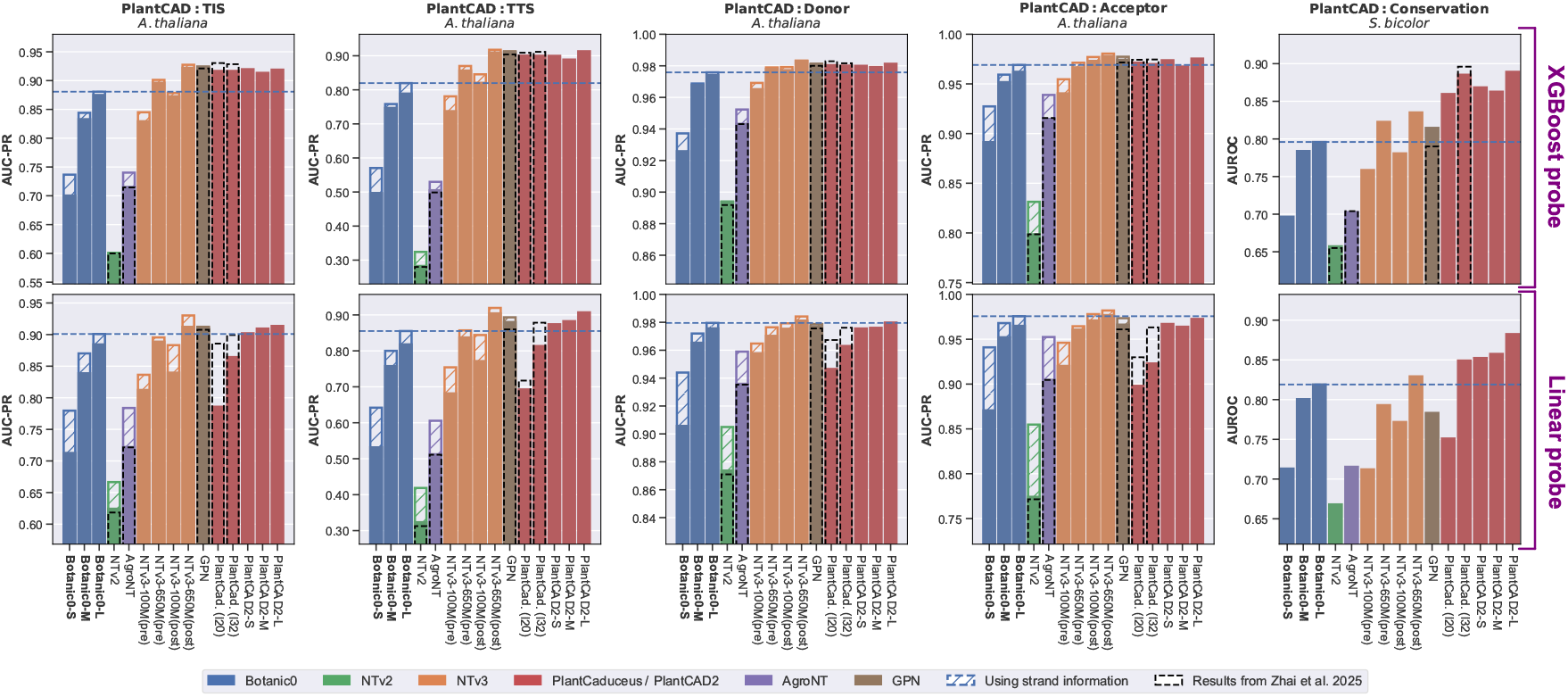
Linear and XGBoost probing results of `Botanic0` models with respect to state-of-the-art models on five binary classification tasks (TIS, TTS, Donor, Acceptor, Conservation) from PlantCAD [24]. Black dashed bars correspond to values reported in [24]. Hashed bars on top of plain-color bars correspond to the increase in performance when using the strand information from the PlantCAD dataset (which is relevant for tasks TIS, TTS, Donor, Acceptor): all negative-strand “-” samples were transformed into their reverse complements. This transformation was applied to all models that are not RC-equivariant (i.e. all models except PlantCaduceus and PlantCAD2 models), and always improved performance. Hashed bars are not visible when the improvement is too small. Better seen in color.

**Table 1:** Comparison of DNA foundation models evaluated in this study.

| Model | # Trainable Parameters | Architecture | Tokenization | Max context length |
| --- | --- | --- | --- | --- |
| AgroNT [1] | 991M | Transformer | 6-mer | 6000bp |
| NTv2-500M [18] | 500M | Transformer | 6-mer | 12kb |
| NTv3-100M [20] | 100M | U-Net + Transformer | Single base | 1Mb |
| NTv3-650M [20] | 650M | U-Net + Transformer | Single base | 1Mb |
| GPN [19] | 66M | CNN | Single base | 512bp |
| PlantCaduceus_L20 [24] | 20M | Mamba | Single base | 512bp |
| PlantCaduceus_L32 [24] | 225M | Mamba | Single base | 512bp |
| PlantCAD2-S [25] | 88M | Mamba-v2 | Single base | 8192bp |
| PlantCAD2-M [25] | 311M | Mamba-v2 | Single base | 8192bp |
| PlantCAD2-L [25] | 694M | Mamba-v2 | Single base | 8192bp |
| **Botanic0-S** | 114M | Transformer | 6-mer | 6000bp |
| **Botanic0-M** | 260M | Transformer | 6-mer | 6000bp |
| **Botanic0-L** | 991M | Transformer | 6-mer | 6000bp |

**Table 2:**
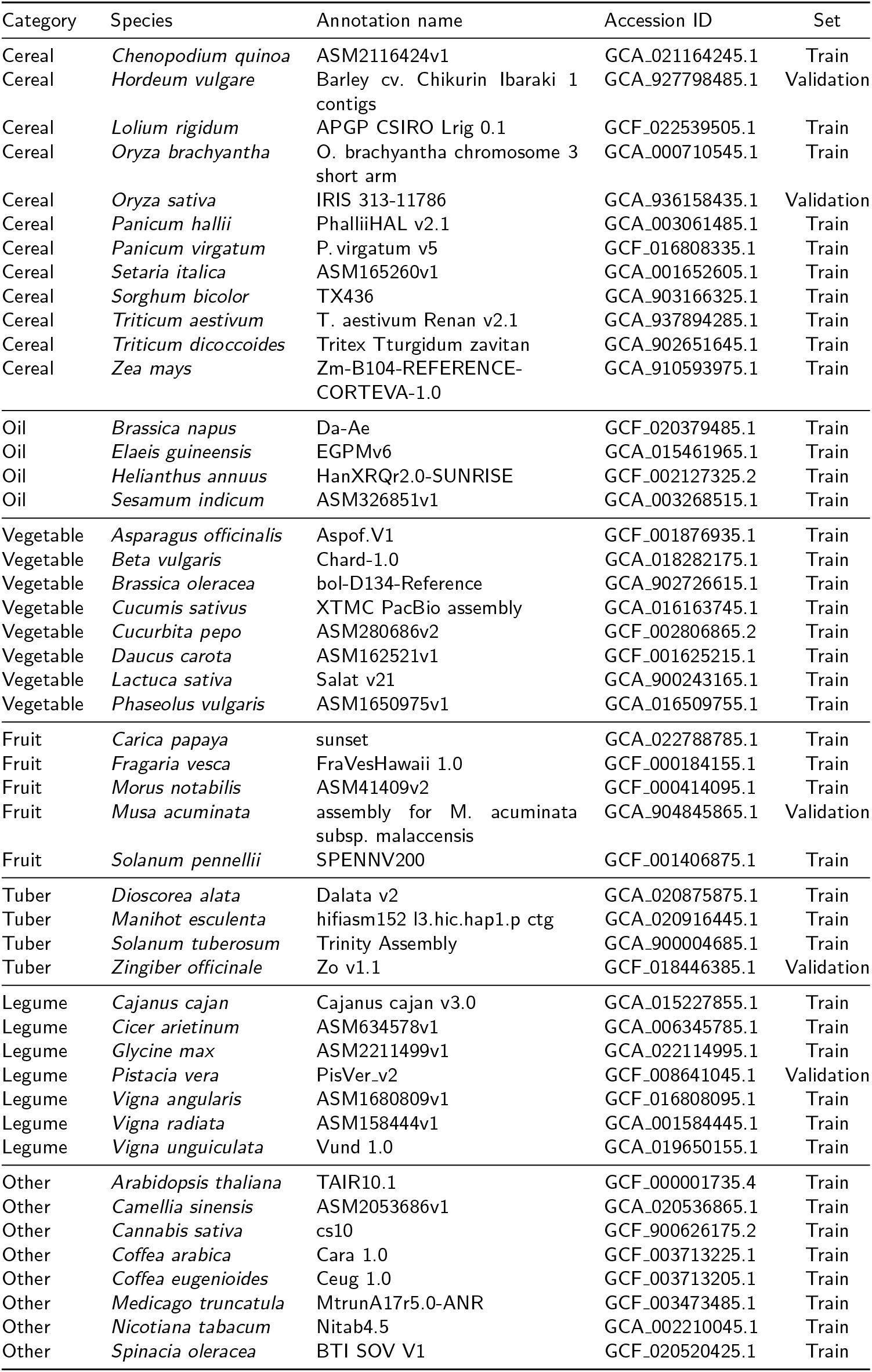
Species and genome assemblies included in the pre-training and validation datasets.

Additionally, we perform a robustness analysis of the `Botanic0` models by varying the *L*2 regularization strength when training the linear probe and show in Supplementary Figure S6 that, unlike PlantCAD [24], the `Botanic0` models are not as sensitive to the regularization strength and showcase a strong performance profile across different regularization strengths. This indicates that in `Botanic0` models, task-relevant information might lie in a well-conditioned, high-variance subspace, allowing a low-norm linear classifier to already capture the signal.

#### 2.3.3 Pre-training longer Botanic0 models leads to better representations

For the five tasks used in Section 2.3.2 as well as the binary classification task “Enhancer Region” from the PGB [1], we further study the performance of `Botanic0` models across their pre-training paths using XGBoost probing. To do so, we performed the same XGBoost probing analysis as in Section 2.3.2 on top of the embeddings generated by different checkpoints of the models. The results, reported in Figure 4, show that most of the time downstream task performance sharply increases over the first ∼60K global steps for all three models. Then performance either plateaus or keeps increasing at a lower rate. In all cases, the performance of `Botanic0` models on downstream tasks as a function of global steps is less smooth and less monotonic than the training and validation loss shown in Figure 1.

**Figure 4.**
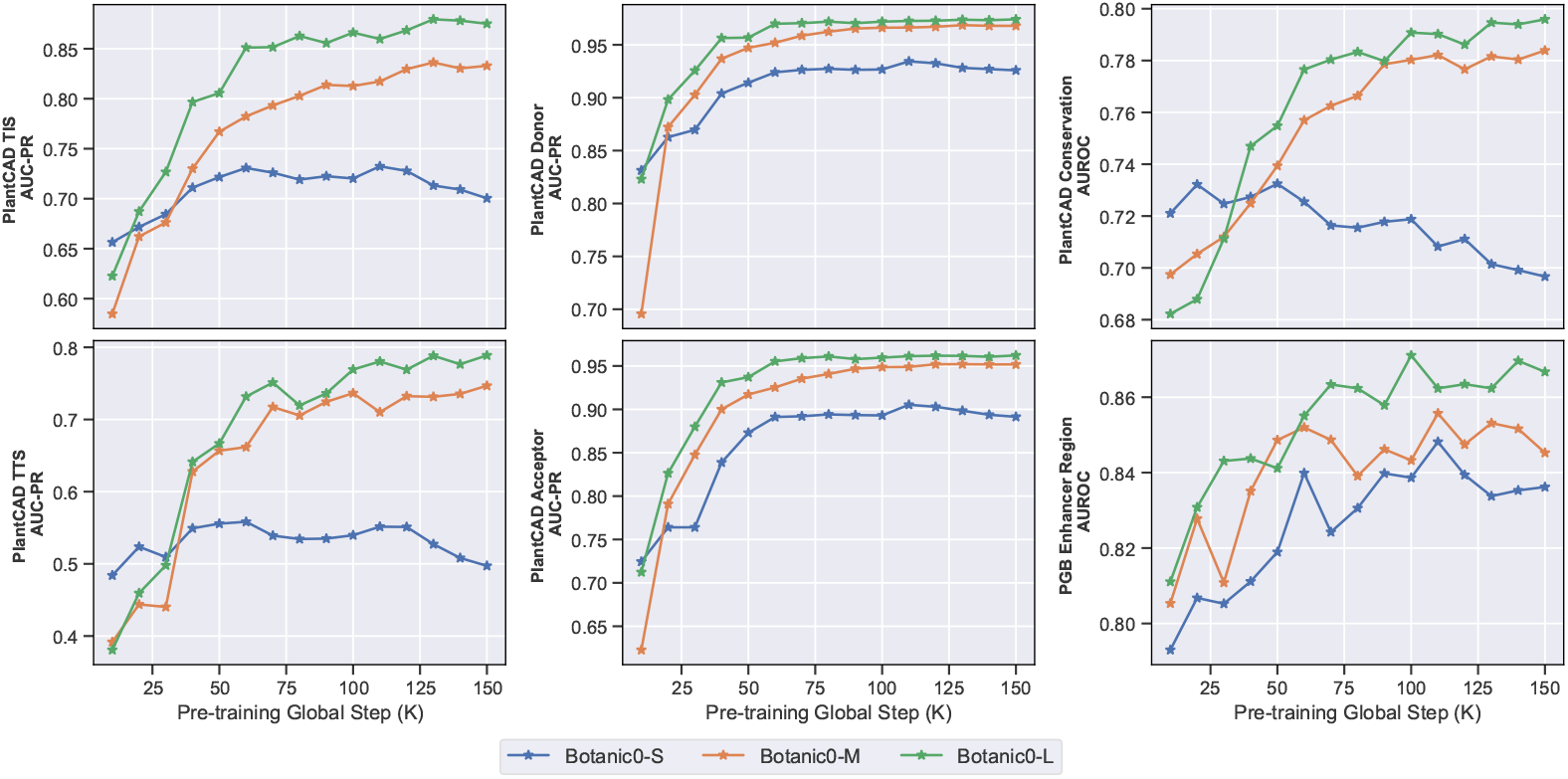
Performance as a function of pre-training steps for each `Botanic0` model using XGBoost probing on six binary classification tasks (cf. Section 4.2.3 for technical details).

### 2.4 Efficient fine-tuning of Botanic0 increases performance on downstream tasks

We further assess the capabilities of the `Botanic0` models by fine-tuning `Botanic0-S`, `Botanic0`-M, and `Botanic0-L` on 17 PGB datasets [1], using IA^3^ [35] following the protocol of [1]. Despite the use of modern GPUs and parameter-efficient fine-tuning (PEFT) methods, fine-tuning remains computationally expensive. For instance, fine-tuning `Botanic0-L` for six epochs on a PGB task for a given hyperparameter configuration takes approximately one hour using an H100 GPU. Consequently, given the number of tasks and models we limited our experiments on the majority of tasks to a small grid search comprising six combinations of learning rates and weight decays, with all other hyperparameters fixed. We observe that, particularly for regression tasks, fine-tuning performance is highly sensitive to hyperparameter choices and optimization noise, complicating direct performance comparisons relative to linear probing. In contrast to [18], which reports greater variance for linear probing than for fine-tuning, we find that linear probing—when performed on a fixed layer—is substantially more robust to hyperparameters and more stable across tasks. Additional details on computational budget constraints, metrics used, grid-search design, and fine-tuning considerations are provided in Section 4.2.4.

We see in Figure 5 the performance of the `Botanic0` family of models and AgroNT on both regression and classification tasks from the PGB. Unlike in the previous section, we see that the gap between `Botanic0` and AgroNT performances narrows indicating that, on such tasks, pre-training is potentially less impactful given enough data. Further evidence for this observation comes from [1], who report that the performance of AgroNT on promoter strength and terminator strength tasks is on par with that of from-scratch baselines from the literature [36, 37].

**Figure 5.**
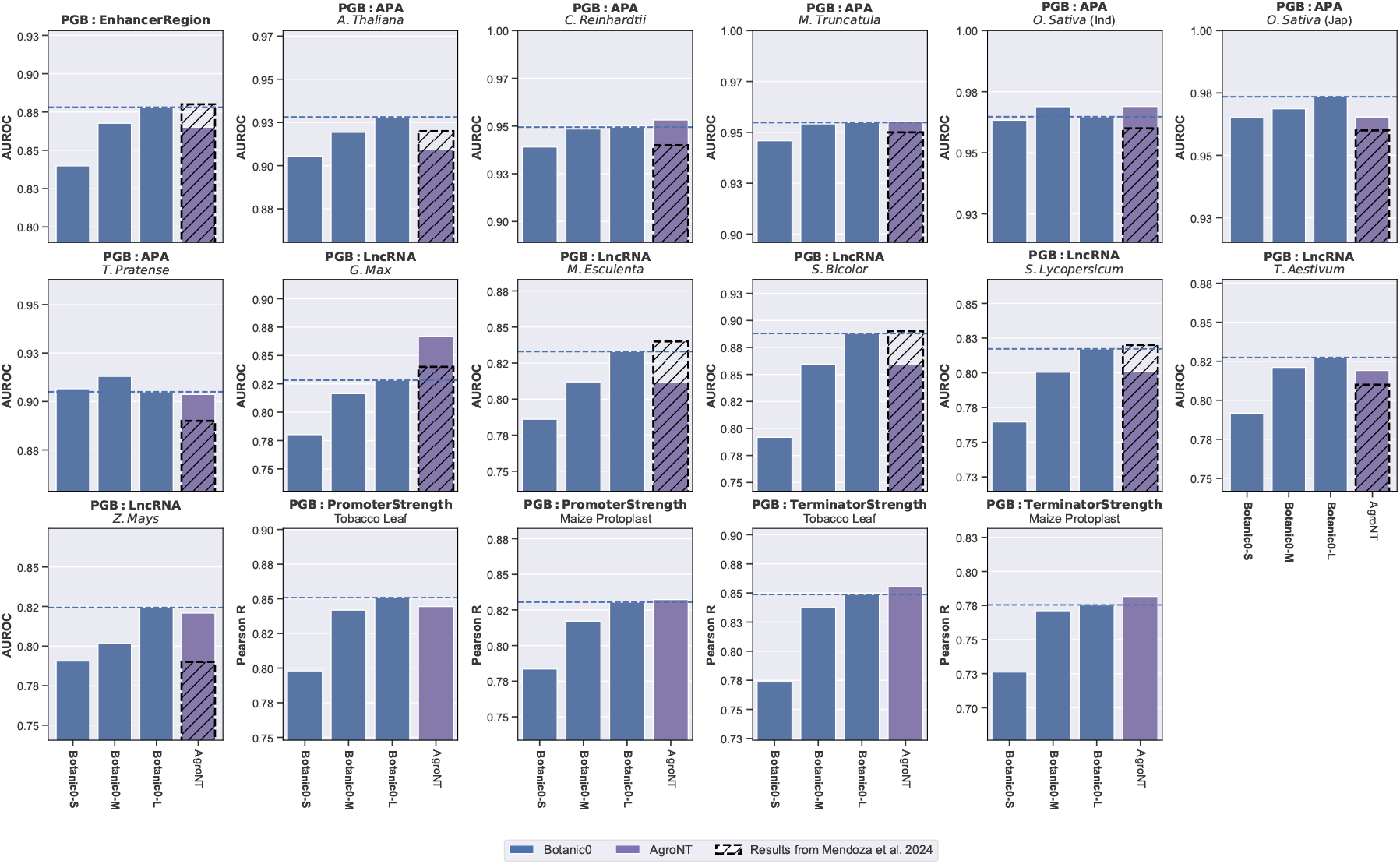
Fine-tuning performance on PGB datasets of `Botanic0` models and AgroNT (cf. Section 4.2.4 for technical details).

## 3 Discussion

This work introduces the initial release of the `Botanic0` model family, intended primarily as a foundation upon which we will build more advanced and refined models. Our pre-training pipeline was inspired by AgroNT [1] although our exact implementation choices differed (see Section 4.1) leading to markedly different foundation models. Our results show that the representational quality of `Botanic0` embeddings, assessed via different means (zero-shot LLR in Sec. 2.2, genomic regions classification in Sec. 2.3.1, probing in Sec. 2.3.2, and fine-tuning in Sec. 2.4), matches or slightly exceeds that of the original AgroNT models, meeting our primary objective.

First, we showed that the MLM loss consistently improved with both model size and training duration, hinting at the existence of “scaling laws” between model performance and computational budget. This relationship would need to be more thoroughly investigated in order to be able to accurately forecast performance at larger scales [38, 39]. Importantly, this favorable scaling extended beyond pretraining metrics: we observed that downstream task performance using frozen model embeddings also improved systematically with increased model capacity and training time.

We also showed that we could fine-tune our models on PGB tasks [1], and obtained results similar to AgroNT. However, it is worth noting that some of these tasks, such as promoter and terminator strength, may not strongly benefit from foundation-model pretraining. Indeed, in many systems, simple sequence statistics — including GC content and the presence of canonical motifs like TATA boxes — correlate with promoter activity and can be useful predictive features in computational models [36, 37]. On such tasks, with sufficient training data, even small deep convolutional neural networks (CNN) models can readily learn these straightforward patterns [36, 37]. We expect the advantages of pretraining to become more pronounced on complex, data-limited tasks, for example, trait- or phenotype-level prediction, where sequence–function relationships are nonlinear, multi-scale, and inherently harder to learn from scratch.

Interestingly, we can already make several observations suggesting immediate directions to explore for improving the performance of the `Botanic0` models in the next versions of the model family. First, we did not observe a performance plateau while scaling the model sizes suggesting that further gains are to be expected with increased model capacity. Second, models built with stronger inductive biases regarding DNA, such as those proposed in [24], which build reverse-complement (RC) equivariance directly into the architecture of the models are promising directions. This is reinforced by our own strand normalization experiment on PlantCAD datasets [24], where models lacking RC-equivariance benefited substantially.

Our findings also contribute to the emerging consensus that single-nucleotide tokenization is preferable for large genomic LMs. Although *k*-mer tokenization has been shown to be suitable for specific tasks at a smaller scale [26], single-base tokenization maximizes resolution, avoids ambiguity in motif representation, and thus appears to be the dominant choice among recent high-performing models. Generalist models such as NTv2 [18], although trained on ≈4, 000 multi-organism genomes (vs. 43 plant-only genomes for `Botanic0`), nonetheless underperform on plant tasks precisely because plants are absent from their training corpus. By contrast, NTv3 [20] retains strong performance on plants despite being trained on a multi-organism dataset, indicating that scale and diversity can partly offset limited domain specificity. However, the efficiency of targeted training remains striking: PlantCAD2 [25], trained on just 65 plant-only genomes, matches or outperforms NTv3 trained on more than 100, 000 full genomes, and our 43-plants model is close to theirs—underscoring that while scale helps, careful in-domain data selection and curation can deliver comparable or better results with orders of magnitude less data. Continuing on this line of thought, given a set of species, it is likely that oversampling the functional regions of the genome in order to balance the ratio of functional regions to intergenic regions (see Section 4.1.1) would allow the model to focus its capacity on learning the functional regions of the genome rather than the intergenic regions. This is what is done in the concurrent works of both Evo2 [23] and PlantCAD2 [25]. Similarly, NTv3 [20] encourages functional regions learning but does it explicitly via functional and regulatory supervision during post-training.

Given the rapid evolution of the LLM architecture landscape [40], adopting more recent architectural design choices [41, 18] is likely to be beneficial. These designs are known to facilitate log-likelihood optimization, and in our benchmark improved likelihood was correlated with stronger performance. It is also interesting to note that the recently published foundation model AlphaGenome [42] was trained on human and mouse DNA using a purely supervised paradigm rather than MLM, highlighting that self-supervision is not the only viable strategy to build genomic foundation models.

More broadly, current benchmarks such as PGB [1] and selected datasets from the PlantCaduceus article [24] curated and made available through Hugging Face have been instrumental for early model development but are inherently limited: they focus on local regulatory signals, motifs, splice sites, promoter strength, rather than integrated organismal traits. These tasks provide valuable insight into mechanistic sequence features but remain far removed from the higher-order processes that determine plant growth, stress tolerance, or yield. Progress toward phenotype-scale prediction will require new benchmarks that bridge this gap by incorporating whole-genome, multi-omic, environmental, and developmental information. Encouragingly, the first signs of this shift are emerging, with recent studies using model-based scores to inform GWAS analyses [24] or to enhance functional variant prediction [19, 43] but current experiments are preliminary and remain limited and qualitative. Ultimately, advancing toward robust trait-prediction models will require moving beyond DNA alone. Plant phenotype is shaped by multimodal interactions: transcriptomics, proteomics, epigenetics, developmental context, environment, and management practices all play essential roles [44]. We envision future generations of Botanic as multimodal foundation models that integrate these diverse data sources to better capture the biological complexity of plant systems and accelerate the development of stress-resilient, high-yielding crop varieties.

## 4 Methods

### 4.1 Pre-training

In this section, we provide a detailed description of the pre-training pipeline used to train the `Botanic0-S`, `Botanic0-M`, and `Botanic0-L` models. Our pipeline is inspired by the approach developed in AgroNT [1]. When implementation details were not specified in the AgroNT paper, we adopted standard choices, which we describe below.

#### 4.1.1 Offline data creation, splitting and processing

Following Suppl. Data 1 of [45], we downloaded 48 assemblies from the National Center for Biotechnology Information (NCBI). We emphasize that we used the assemblies from the supplementary materials of the v1 bioRxiv preprint version of the AgroNT paper [45], which are different from the assemblies listed in the published version of the same paper [1]. For convenience, we provide the full list of assemblies we used in the pre-training in Supplementary Table 2.

We display in Figure 6 a high-level overview of the total genomic content of the pre-training and validation datasets. In this figure, to facilitate visualization we used the updated assemblies from the published version of the AgroNT paper [1] as they all have available genome regions annotations in contrast to the assemblies we used in the pre-training pipeline for which annotations are more scarce. As a comparison point, we add in Supplementary Figure S8 the same plot but with the original assemblies we used in the pre-training pipeline where we labeled entire unannotated genomes as “intergenic”.

**Figure 6.**
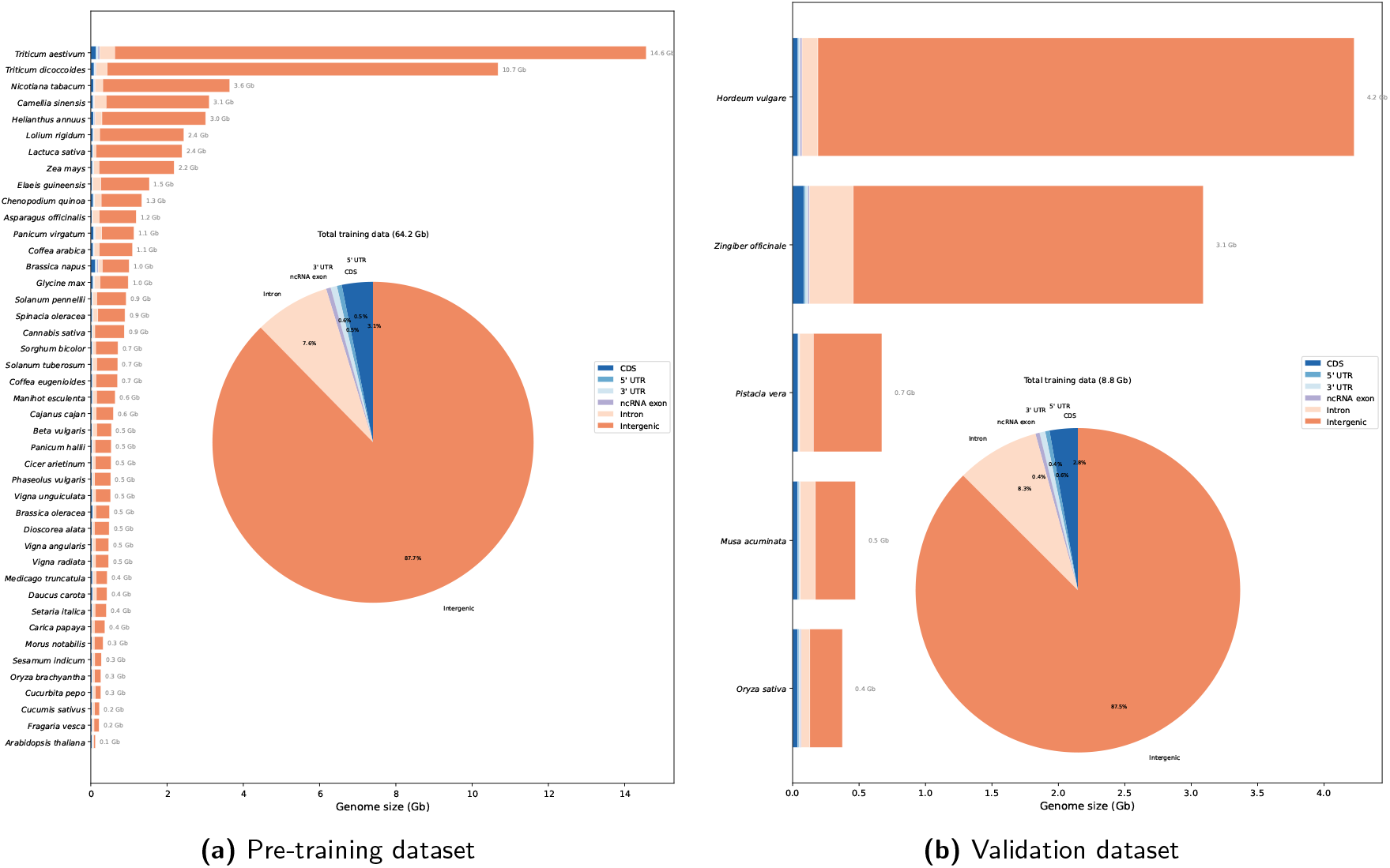
Overview of the genomic content of the pre-training and validation datasets using the updated assemblies from the published version of the AgroNT paper [1].

For each assembly, we used the full-genome FASTA file rather than individual chromosome files, retaining only nuclear DNA (i.e., excluding mitochondrial, chloroplast, or other plastid DNA) and filtering out entries shorter than 6,100 bp - this includes chromosomes, contigs, and scaffolds that passed the length threshold. We then split each retained sequence from start to end into overlapping windows of 6,100 bp with a 50 bp overlap. It is apparent from Figure 6 that with uniform sampling of the genome as we do here, the pre-training dataset is dominated by intergenic regions reaching more than 85% of the genomic content. The resulting dataset contains 11,488,516 sequences, stored in jsonl format which is better suited to cloud-based streaming setup compared to fasta files.

Out of the 48 species downloaded, we used 43 of them as the training dataset, and 5 species were used as a hold-out validation dataset. This validation dataset corresponds roughly to 10% of the overall data, and contained the following species: *Oryza sativa* (`IRIS 313-11786`), *Pistacia vera cultivar Batoury* (`PisVer_v2`), *Hordeum vulgare subsp. vulgare* (`Barley cv. Chikurin Ibaraki`), *Zingiber officinale cultivar Zhangliang* (`Zo_v1.1`), *Musa acuminata subsp. malaccensis* strain Doubled-haploid Pahang (`GCA_904845865.1`). Thus, these 5 species are never seen by the models during pre-training. In terms of number of sequences this amounts to keeping 10, 026, 502 sequences of length 6, 100 bp for training and 1, 462, 014 sequences of length 6, 100 bp for validation. In the rest of the article, we will refer to these datasets as the pre-training and validation datasets and consider each sequence of size 6, 000 bp as a sample.

As a final preprocessing step, we loaded both datasets in RAM once and shuffled the pre-training (and validation) datasets independently to ensure that the order of the batches of sequences when training the models was sufficiently random as we only use approximate shuffling techniques at training time (see Section 4.1.4). This is an important step as we noticed that, in the absence of shuffling when sequences were ordered by species and chromosomes, the training process was considerably degraded.

#### 4.1.2 Online tokenization, data augmentation and masking

Our pre-training implementation is based on the lingua library [46], which we forked and adapted to both DNA data and masked language modeling (MLM). In particular we added support for the n_views=1 (single view) case and added random states to the tokenizer. This enables extracting, in an online fashion, a random window of size 6, 000 bp from the 6, 100 bp long sequences as in [1] and applying random masking, while keeping the perfect reproducibility and resumability of the training data generation process, which was one of the main features that directed us to use the lingua library [46].

We implemented a 6-mer tokenization process similar to [1], where we fixed the length of our output sequences to 1, 024 tokens (including the special `<CLS>` token) instead of 1, 025 for GPU computational efficiency reasons (GPUs break down computations into warps which are multiples of 32 threads). We then used the same masking hyper-parameters as in the original BERT paper [10].

#### 4.1.3. Model Training

Our models follow the exact architecture described in [1], which corresponds to an ESM-2 model [47] repurposed for DNA data. Our three models (`Botanic0-S`, `Botanic0-M`, and `Botanic0-L`) have respectively 4, 10, and 40 layers and `Botanic0-L` matches the parameters count of AgroNT [1]. We use the traditional cross-entropy loss function over the masked tokens for the loss computation.

For the initialization scheme, we used the `current_depth` scheme [48] from lingua, which scales the initialization of the weights by a factor of 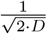 where *D* is the depth of the layer. Additionally, we make sure the input embedding, which is the sum of the learned positional embeddings and token embeddings, is normalized to have a mean of 0 and a standard deviation of 1.

Our models were trained using AdamW optimizer [49] with the same learning rate schedule as in [1], which is a special case of the well known warmup-stable decay [50] learning rate schedule with a warmup period of 64, 000 steps, no stable period and an inverse square root decay function with a 315, 000 steps horizon. A “step” is defined as the processing of exactly `grad_acc · num_gpus · batch_size` sequences (or samples) of size 6, 000 bp (or 1, 024 tokens) or equivalently `grad_acc · num_gpus · batch_size · num_tokens_per_sequence` tokens. Where `grad_acc` is the gradient accumulation factor, `num_gpus` is the number of GPUs, `num_tokens_per_sequence` is the number of tokens per sequence and `batch_size` is the number of sequences per worker’s batch (or samples per batch per worker).

For instance for the `Botanic0-L` model we used `grad_acc=23, num_gpus=8`, `num_tokens_per_sequence=1024` and `batch_size=8`, thus processing an equivalent batch of samples of size of 8 · 8 · 1024 · 23 = 1, 507, 328 tokens (we similarly increase batch sizes for smaller models, while keeping the total number of samples per batch constant by reducing the gradient accumulation factor thus speeding up the training process considerably). This is only an upper bound on the true batch size, as only masked tokens are considered for the loss computation; only roughly 15% of the tokens are actually used for the loss computation. Thus the actual batch size is only ≈ 226, 100 tokens.

We trained our models for approximately 150, 000 steps instead of 364, 000 steps as reported in [1], as we found that training longer is both costly and produces only marginal improvements in terms of final loss, even though we found that performance on downstream tasks was not yet saturated for the `Botanic0-L` model.

#### 4.1.4 Hardware and system considerations

We trained the models using `bfloat16` precision [51] on different clusters of machines with different hardware configurations but using each time 8 GPUs, whether they were *A*100, *H*100 or *B*200 GPUs. Some of our trainings were scheduled using the Slurm-based job scheduling [52].

We split the two jsonl files containing the pre-training and validation datasets into 8 chunks of roughly equal size in order to increase the efficiency of the distributed data parallel (DDP) training, which is always done on a multiple of 8 GPUs.

Finally, we use a strategy similar to reservoir sampling [53] for training-time shuffling as implemented in the lingua library [46]. We apply this sampling technique on each worker with a prefetch size parameter (counted in terms of batches) in order to be able to approximately shuffle the data while keeping RAM consumption under control.

With this setup, the training of the largest model, `Botanic0-L`, took roughly 10 days to complete on a single *B*200 GPU node (with 8 *B*200 GPUs).

#### 4.1.5 Genomic region classification

Following the GPN [19] methodology, we tiled the *Arabidopsis thaliana* (TAIR10) nuclear genome into non-overlapping 100 bp windows using FASTA and GTF files from Ensembl Plants release 59 [54], and repeat annotations from the UCSC Table Browser^‡^. Each window was assigned one of seven labels—`cds`, `utr3, utr5, intron, ncrna, intergene`, or repeat—requiring 100% coverage by the corresponding annotation. Introns were computed as protein-coding gene regions minus exons; intergenic windows (zero gene overlap) were further split into `repeat` (fully covered by repeat elements from a BED file) and `intergene`. Ambiguous windows matching multiple labels were discarded. For each labeled window, a 512 bp context sequence centered on the 100 bp region was extracted and passed through the model; the token embeddings covering the central 100 bp were then mean-pooled to obtain a single per-window representation. We note that 512 bp is suboptimal for `Botanic0` as it was trained on 6000 bp windows, but we keep 512 for simplicity. Finally, we add that, because of the 6-mer tokenization and the presence of “N”, which are tokenized as 1-mer, in some sequences identifying the token embeddings representing the central 100 bp region is non-trivial and introduces some overhead. These embeddings were used both as inputs to pymde [33] for minimum-distortion embedding visualization, and for multi-class classification with XGBoost [32], holding out chromosome 5 as the test set.

#### 4.1.6 Zero-shot variant scoring via log-likelihood ratios

To evaluate whether the pre-trained models capture functional constraints in the genome, we computed zero-shot log-likelihood ratios (LLR) for single-nucleotide variants in *Arabidopsis thaliana*. The LLR of a variant is defined as the difference in log-probability assigned by the model to the alternative allele versus the reference allele at a masked position:

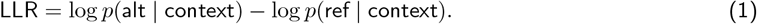

A negative LLR indicates that the model considers the alternative allele less likely than the reference, consistent with functional constraint at that position [15]. We expand on the details of the masking procedure below in Sec. 4.1.6.2.

##### 4.1.6.1 Variant categories

We constructed a benchmark of ∼ 100 variants per category from the TAIR10 genome annotations (GTF), targeting positions where a single-nucleotide change is expected to be deleterious or neutral:

- **Start codon**: first nucleotide of annotated ATG start codons (A→*{*C,G,T*}*);
- **TATA box**: position 3 of TATAAA motifs found 25–35 bp upstream of 5^*′*^ UTR starts (A→*{*C,G,T*}*);
- **Splice donor**: second nucleotide of GT donor dinucleotides at intron boundaries (T→*{*A,C,G*}*);
- **Splice acceptor**: first nucleotide of AG acceptor dinucleotides (A→*{*C,G,T*}*);
- **PolyA signal**: position 3 of AATAAA motifs in 3^*′*^ UTRs (A→*{*C,G,T*}*);
- **CDS codon positions**: random substitutions at each of the three codon positions in verified protein-coding exons, with all three possible alternative alleles emitted per site (*∼*100 sites *×* 3 alternatives per position);
- **Neutral (intronic)**: random substitutions at interior intronic positions (≥10 bp from splice sites);
- **Neutral (intergenic)**: random substitutions in intergenic regions ≥5 kb from any annotated gene.

Only canonical transcripts on the (+) strand were considered for extracting functional annotations. Introns were inferred from consecutive exon boundaries within each transcript. Intergenic gaps shorter than 10 kb were excluded to avoid proximity to genic regions.

##### 4.1.6.2 Scoring procedure

Each variant was embedded in a 6,000 bp genomic context window centered on the variant position. For `Botanic0` and AgroNT, the 6-mer containing the variant nucleotide was masked and the LLR was computed from the logits of the reference versus alternative 6-mer tokens. For single-nucleotide–tokenized models (GPN, PlantCaduceus), the variant nucleotide itself was masked and scored directly. GPN scores were averaged over forward and reverse-complement strands to account for its non–RC-equivariant architecture; PlantCaduceus is natively RC-equivariant and was scored on the forward strand only, using its native 512 bp context window.

### 4.2 Probing and Fine-tuning

#### 4.2.1 Datasets

We used two sets of datasets for downstream tasks in our experiments. The first one is the Plant Genome Benchmark (PGB) from [1], which contains 8 tasks including regression (promoter strength and terminator strength), binary classification, and multi-label classification tasks. These datasets are available on HuggingFace under the repo name `InstaDeepAI/plant-genomic-benchmark`. We also used five datasets from [24] available on HuggingFace under the repo name `kuleshov-group/cross-species-single-nucleotide-annotation`. These datasets correspond to the binary classification translation initiation sites (TIS), translation termination sites (TTS), splice donors and acceptors and conserved sites. We refer to them as PlantCAD-TIS, PlantCAD-TTS, PlantCAD-Donor, PlantCAD-Acceptor, and PlantCAD-Conservation in our work. Four of these datasets (all but PlantCAD-Conservation) are highly imbalanced with 2-5% positive samples in their train and test datasets. Therefore, we use AUC-PR instead of AUROC as evaluation metrics on these datasets. These four PlantCAD datasets also contain samples from both DNA strands each marked as (+) and (*−*) and mixed together.

#### 4.2.2 Competitor models

In Section 2.4, we compared our models to other recent DNA foundation models from the literature. All evaluated models are detailed in Table 1 and were downloaded from HuggingFace (HF) repositories. More precisely, `PlantCaduceus-20`, `PlantCaduceus-32` [24] were downloaded from HF repositories `kuleshov-group/Plant Caduceus_l20` and `kuleshov-group/PlantCaduceus l32`, PlantCAD2-S, PlantCAD2-M, PlantCAD2-L [25] were taken from kuleshov-group/PlantCAD2-Small-l24-d076, `kuleshov-group/PlantCAD2-Medium-l4 8-d1024`, and `kuleshov-group/PlantCAD2-Large-l48-d1536`. Similarly, `NTV3-100M-pre, NTV3-650M-pre, NTV3-100M-post`, and `NTV3-650M-post` [20] were taken from HF repositories InstaDeepAI/NTv3 100M pre, In staDeepAI/NTv3 650M pre, InstaDeepAI/NTv3 100M post, and `InstaDeepAI/NTv3 650M post` respectively. Finally, GPN was taken from HF repository `songlab/gpn-brassicales` [19].

#### 4.2.3 Probing

We use two probing methods in this work in Figures 3 and 4: a linear classifier probe and a XGBoost classifier probe. XGBoost model training was performed using the xgboost library [32]. In order to be consistent with experiments from [24] we used the following hyperparameters: `learning rate=0.1,depth=6,nb epochs=1000`. For the other main hyperparameters we used l2 reg=1.0,l1 reg=0. For the PGB-Enhancer tasks, we decrease the learning rate to 0.05 in all XGBoost experiments as training curves were too unstable otherwise. For the linear classifiers, we used the LogisticRegression implementation from scikit-learn [55] with `C=1.0,l1 reg=0.0,prob e tol=1e-4,fit intercept=True,intercept scaling=1,solver=liblinear,max iter=1000,warm star t=True`.

XGBoost and Linear probings were applied on the embedding of the “middle” token for PlantCAD tasks and on the mean of embeddings of all the tokens for PGB-Enhancer tasks (cf. 4.2.3.1).

For all datasets we precompute embeddings for all sequences once and for all. Regarding PlantCAD datasets we also compute embeddings for a version of each task where each strand was converted to (+) strand representation by reverse-complementing each (*−*) strand.

##### 4.2.3.1 Model specific considerations

We emphasize that, for each model, we probe the “default” embedding layer of the model: the one exposed by the model implementation. We do not grid-search over the layers of the models to select the best one for the fine-tuning task as is done in [18]. [18] has already demonstrated that the best layer for a given fine-tuning task is often not the default one and thus the performance of all models would be improved by fine-tuning or probing different task-specific layers.

Regarding the embedding layer of NTv3 models [20] that we target, we use the nucleotide-level embedding from the “deconvolutional” tower and not the “transformer” tower [20] embedding which is consistent with how [20] performed their fine-tuning experiments.

In addition, among the NTv3 models, `NTV3-650M-post` and `NTV3-100M-post` require, in addition to the input tokens, to pass the name of the considered species as a string parameter to the model. The supported species are the following: *Amphiprion ocellaris, Arabidopsis thaliana, Bison bison bison, Caenorhabditis elegans, Canis lupus familiaris, Chinchilla lanigera, Ciona intestinalis, Danio rerio, Drosophila melanogaster, Felis catus, Gallus gallus, Glycine max, Gorilla gorilla, Gossypium hirsutum, Human, Macaca nemestrina, Mouse, Oryza sativa, Rattus norvegicus, Salmo trutta, Serinus canaria, Tetraodon nigroviridis, Triticum aestivum, Zea mays*. When the species of a sample from a given task was not in this list, we used *Arabidopsis thaliana*. For PlantCaduceus models [24, 25] we additionally use the same reverse-complement (RC) flipping and averaging scheme to produce RC-equivariant embeddings as done in [24] for probing and RC-equivariant scoring by averaging logits over both strands embeddings. We refer to the model code from [24, 25] for more details.

Finally for PlantCaduceus models we use, as a sequence embedding, in the absence of `<CLS>` token, the embedding of the token localized in the middle of the sequence for PlantCAD-TIS, PlantCAD-TTS, PlantCAD-Donor and PlantCAD-Acceptor datasets. Each sequence of these datasets is 512 nucleotides long, so we used the token corresponding to the nucleotide 255 for the models using token at nucleotide level, and the token 43rd for the models using 6-mer. We stress that this construction is only approximate for 6-mer models as N nucleotides, which are tokenized as 1-mer (`<N>`), make the concept of “middle” token somewhat ill-defined even for a given sequence length (this is the exact same problem we already mentioned in Sec. 4.1.5). For the PGB, Enhancer region task [1], we considered the mean of the embeddings of all tokens, as this led consistently to better results for all models.

#### 4.2.4 Fine-tuning

We test the IA^3^ training technique [35] to fine-tune the `Botanic0` models as in [1] on all transformer layers and add a two-layer MLP classifier head with a bottleneck dimension of 1500 (the same as the input embedding dimension) and tanh activations on top of the model’s `<CLS>` token. As most free parameters are located in the classifier head (roughly 2.3*M* parameters irrespective of the model size depending on the target dimensionality), for a target of dimension 1, this amounts to training a total of respectively 1.93%, 0.88% and 0.26% of the total parameters of the `Botanic0-S`, `Botanic0-M`, and `Botanic0-L` models, leaving all other parameters frozen.

For IA^3^ optimization we use the AdamW optimizer [49] with a fixed batch size of 16 and perform two versions of the same grid search: a coarse and a fine-grained grid search.

- **Coarse grid search:** `learning_rate:` [5 *×* 10^*−*3^, 2.5 × 10^*−*3^] and `weight_decay`: [1 *×* 10^*−*5^, 1 × 10^*−*4^, 1 × 10^*−*3^], totaling 6 different fine-tuning combinations.
- **Fine-grained grid search:** `learning_rate:` [10^*−*4^, 2.5 × 10^*−*4^, 5 × 10^*−*4^, 10^*−*3^, 2.5 *×* 10^*−*3^, 5 × 10^*−*3^] and `weight_decay`: [10^*−*5^, 10^*−*4^, 10^*−*3^, 10^*−*2^], totaling 24 different fine-tuning combinations for regression tasks.

The coarse grid search is performed for all tasks and the fine-grained grid search is performed only for regression tasks.

We use warm-start as well as a linear decay learning rate schedule and standard early stopping with a patience of 3 epochs.

We also test LoRA [56] in parallel, a similar, more widely used PEFT method, which is now the recommended method to fine-tune transformers from the NT family [57] but we get roughly similar results as IA^3^ and thus keep IA^3^ as our preferred method to ease comparison with [1]. We show some of the results of LoRA on the PGB regression tasks in Supplementary Figure S5, also illustrating the high variability of fine-tuning results when using different hyperparameters or techniques.

As we are performing a grid search we set aside 15% of the training data as a validation set and use it to find the best hyperparameters; we then retrain the model on the full training set with the best hyperparameters and compute test performance on the test set.

Regarding regression tasks we use Pearson correlation as it is less sensitive to small differences in the prediction values than *R*^2^ score. We note that, irrespective of the pretrained backbone used (AgroNT or `Botanic0`), for regression tasks we obtain lower performance than reported for AgroNT in [1], indicating that the classifier head we used in our experiments and/or our choice of hyperparameters are not optimal for those specific regression tasks.

## 5 Acknowledgements

We thank Benjamin Trom for helpful discussions about pre-training best practices as well as Mathieu Andreux for technical advice. We thank Jean-François Reboud for his help setting up distributed training. We thank Gregory Andrews for his insights on pretraining data composition and his help making Figure 6 and Supplementary Figure S8. Finally, we thank Adnane Boualem, Gregory Andrews, Bertrand Gakière, Benjamin Trom, Félix Raimundo, Julián García-Abadillo Velasco and Diego Jarquín for proofreading the manuscript and providing valuable feedback.

## A Supplementary Material

**Supplementary Figure S1.**
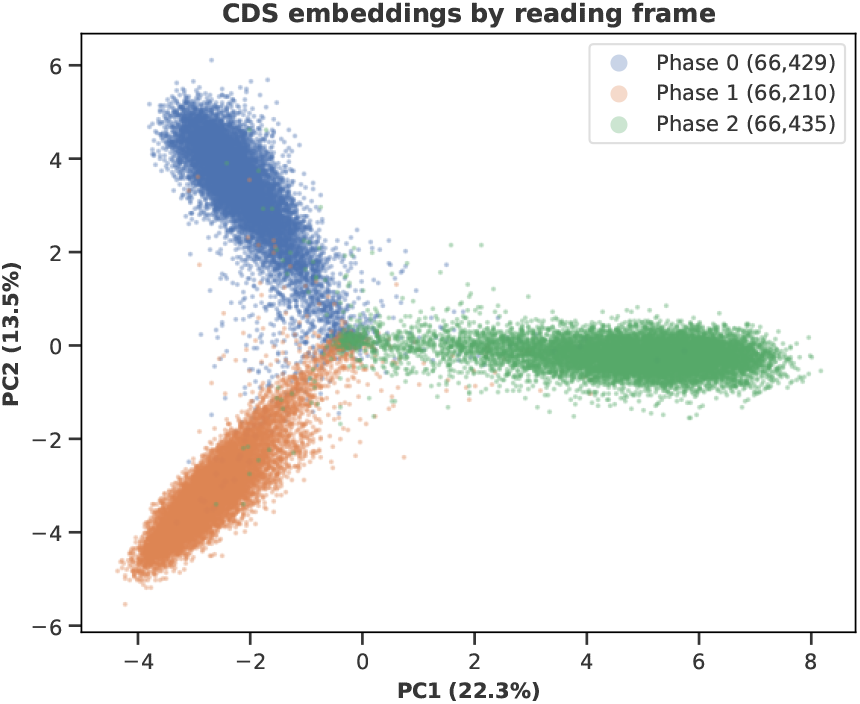
PCA of `Botanic0-L` mean-pooled embeddings for the 199,074 windows labeled as CDS in the *A. thaliana* genomic region classification task (see Section 4.1.5). Each 100 bp window is colored by its strand-aware codon reading frame phase (0, 1, or 2), determined from the GTF CDS frame and strand columns.

**Supplementary Figure S2.**
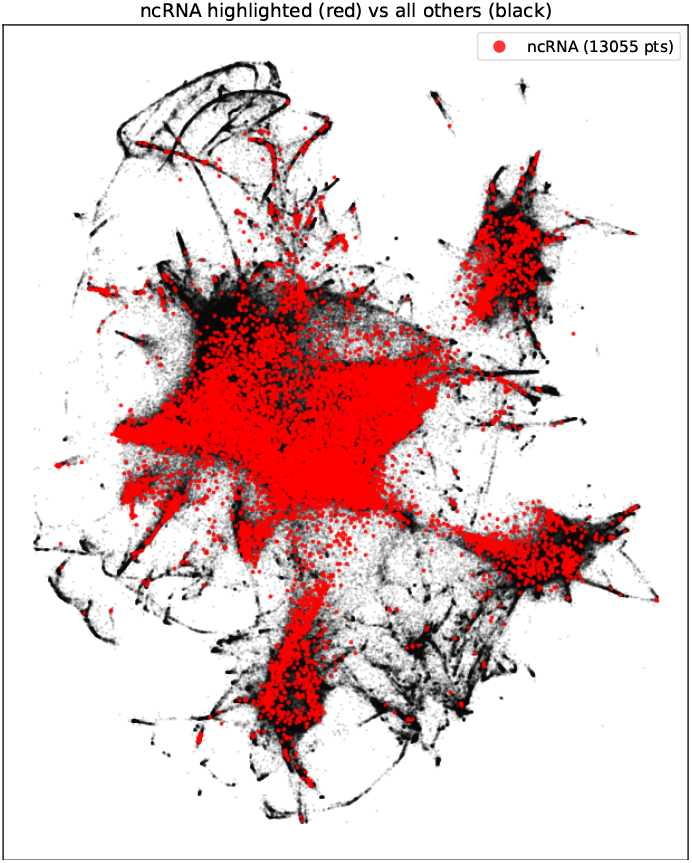
Minimum-distortion embedding of ncRNA class vs the other classes.

**Supplementary Figure S3.**
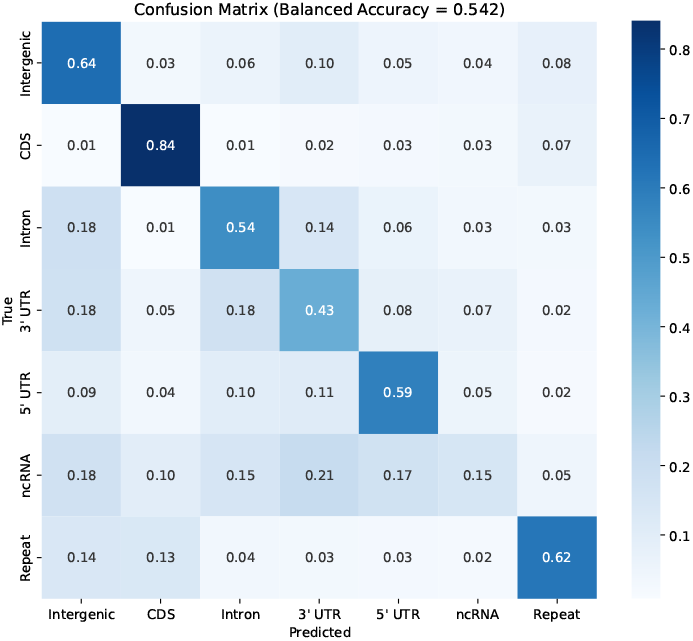
Confusion matrix of the XGBoost classifier for genomic region classification in *Arabidopsis thaliana* from AgroNT embeddings.

**Supplementary Figure S4.**
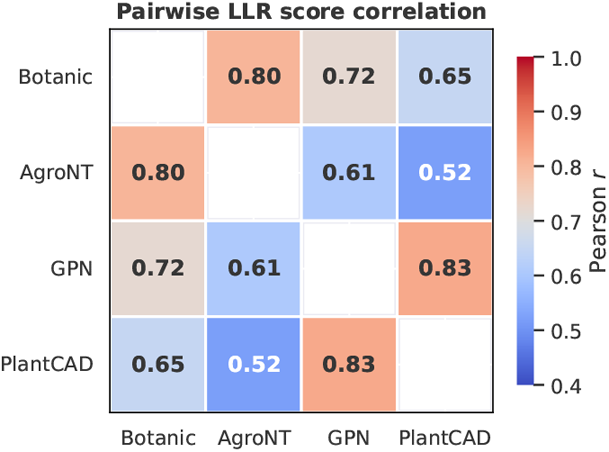
LLR comparison between `Botanic0` models, PlantCAD and GPN models (see Section 4.1.6).

**Supplementary Figure S5.**
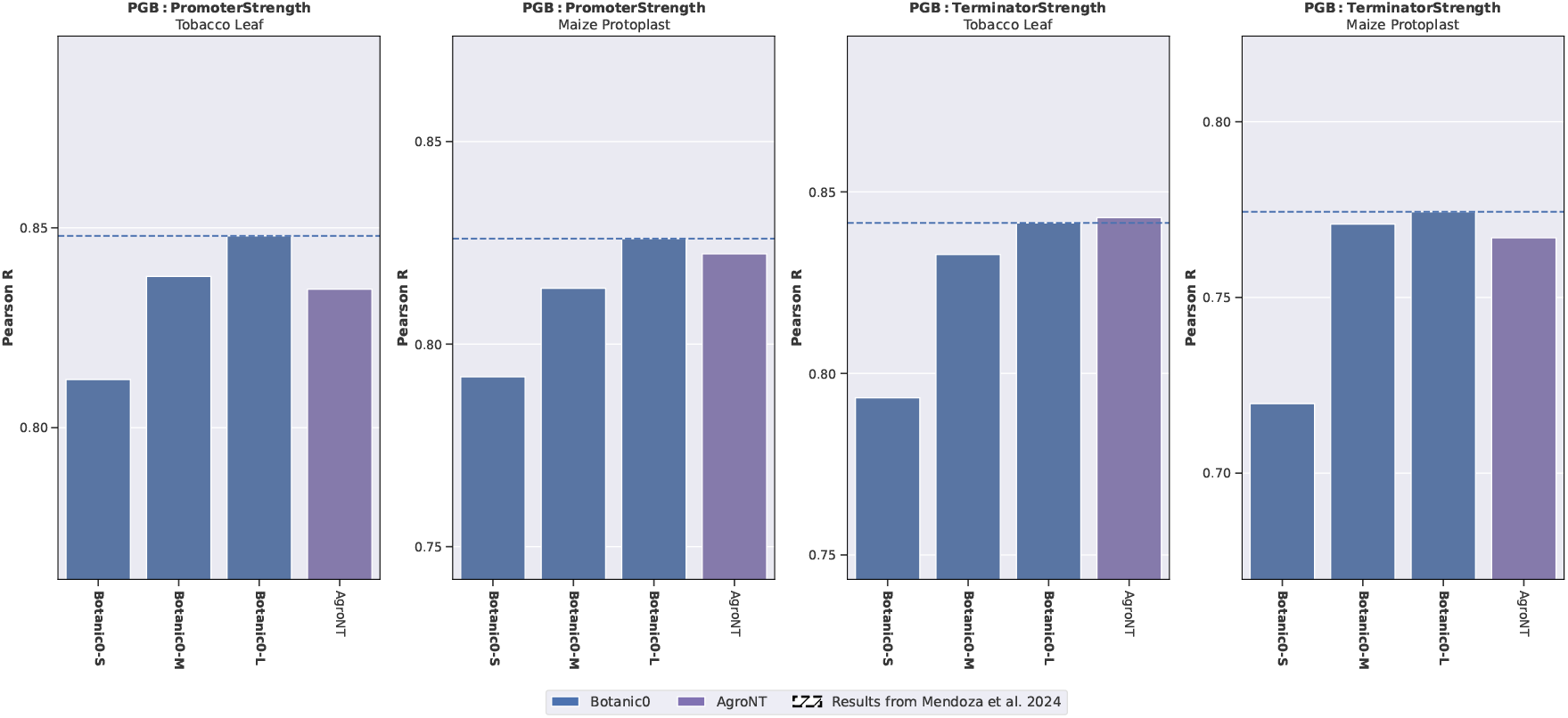
Fine-tuning performance of LoRA on PGB regression datasets of `Botanic0` models and AgroNT.

**Supplementary Figure S6.**
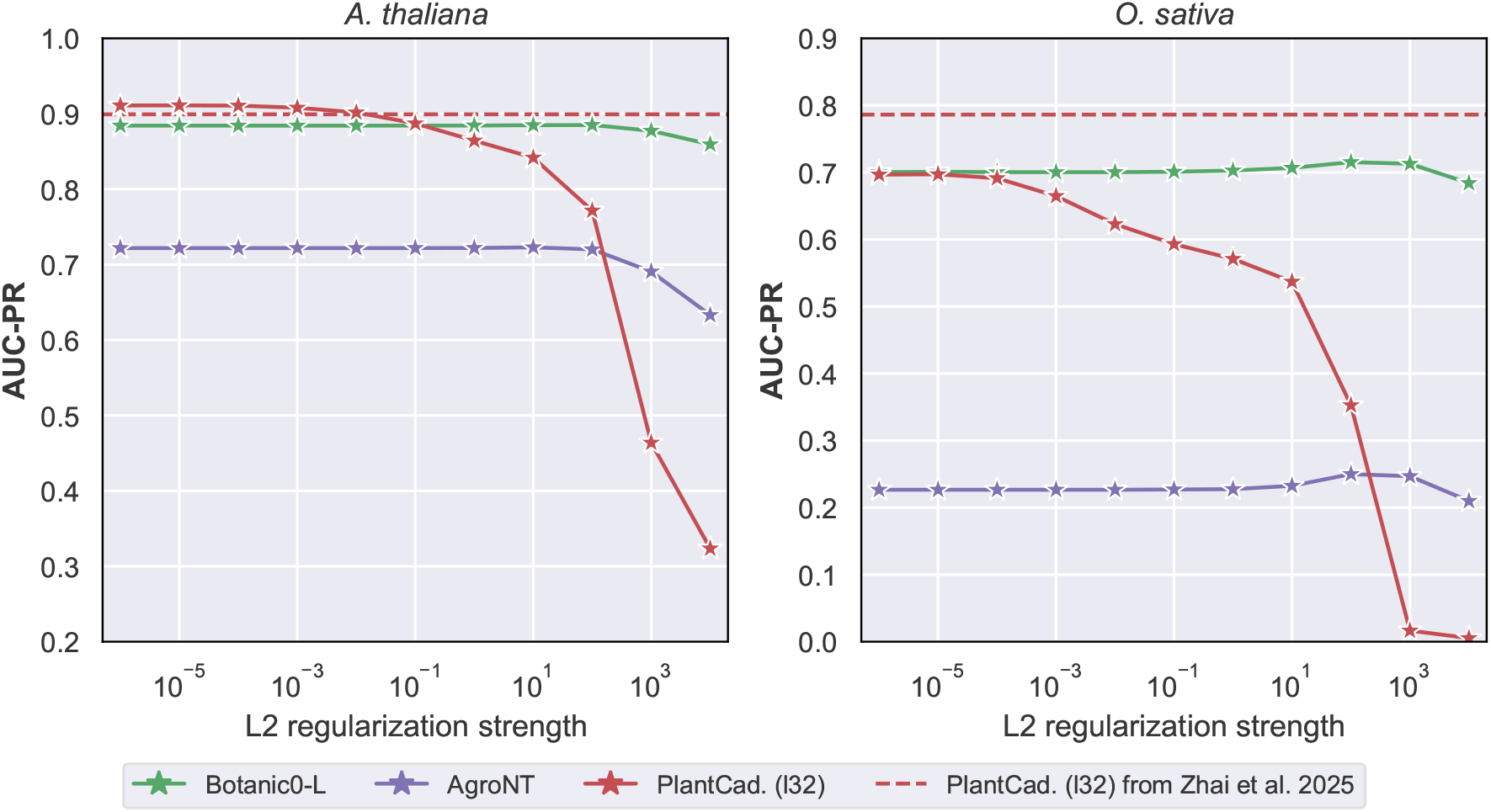
Performance of linear models in PlantCAD-TIS task, trained on chr 1-4 of *Arabidopsis thaliana* and tested either on chr. 5 of *Arabidopsis thaliana* or on *Oryza sativa*.

**Supplementary Figure S7.**
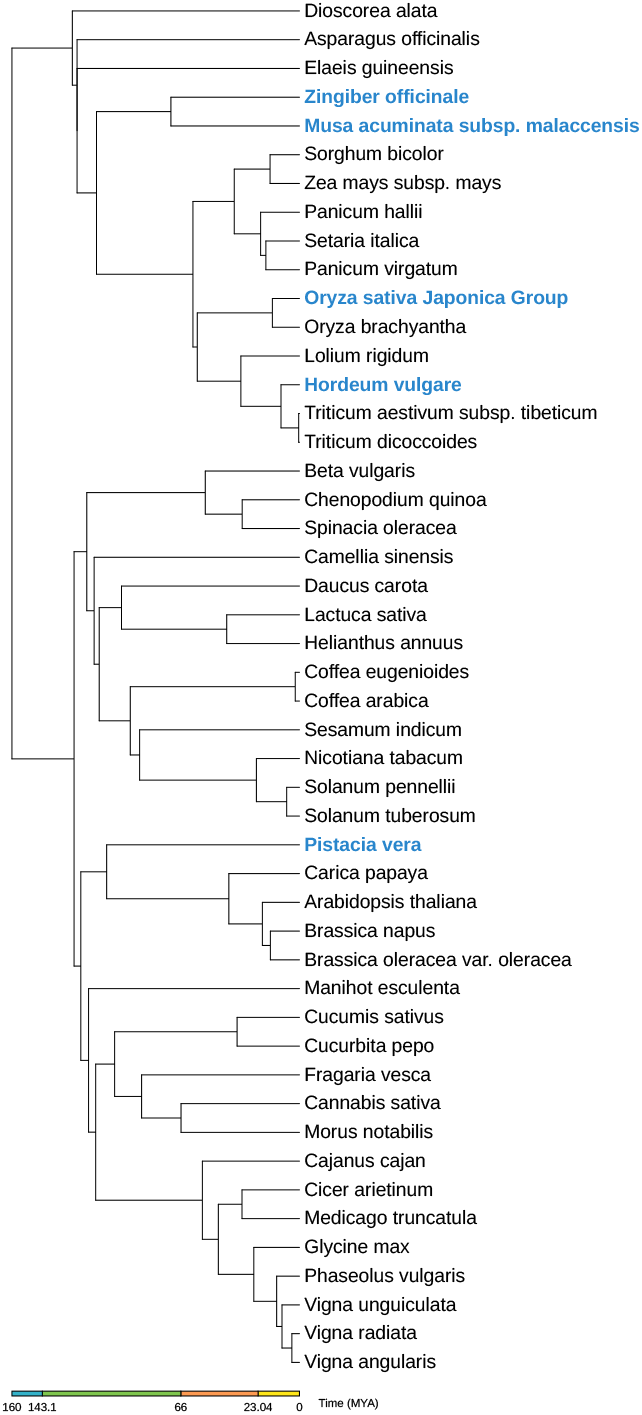
Phylogenetic tree of the species used in the pre-training dataset. The species shown in bold blue are those used for the validation set. The tree was constructed using TimeTree 5 [58] and displayed using iTOL [59].

**Supplementary Figure S8.**
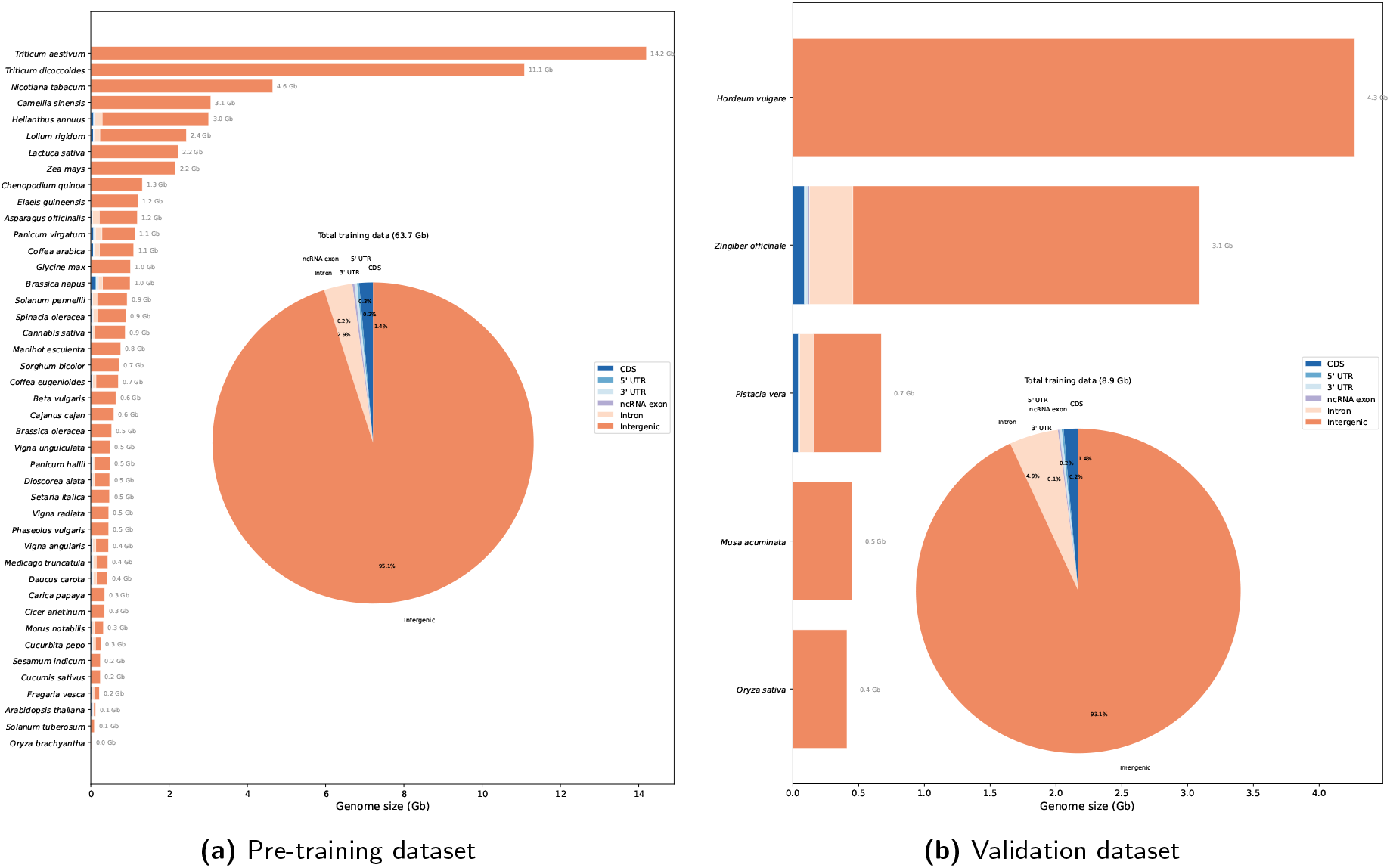
Overview of the genomic content of the pre-training and validation datasets.

## Footnotes

¶ Biological Omics Transformer for Agricultural and Nutritional trait Inference in Crops

‡ https://ucsc.gao-lab.org/cgi-bin/hgTables

